# Exercise performance is not improved in mice with skeletal muscle deletion of natriuretic peptide clearance receptor

**DOI:** 10.1101/2022.08.30.505918

**Authors:** Brigitte Jia, Alexander Hasse, Fubiao Shi, Sheila Collins

## Abstract

Natriuretic peptides (NP), including atrial, brain, and C-type natriuretic peptides (ANP, BNP, and CNP), play essential roles in regulating blood pressure, cardiovascular homeostasis, and systemic metabolism. One of the major metabolic effects of NP is manifested by their capacity to stimulate lipolysis and the thermogenesis gene program in adipocytes, however, their metabolic effect on skeletal muscle is much less appreciated. There are three NP receptors (NPR): NPRA, NPRB, and NPRC, and all three NPR genes are expressed in C2C12 myocytes. Treatment with either ANP, BNP, or CNP evokes the cGMP signaling process in C2C12 myocytes. We then generated a genetic model with *Nprc* gene deletion in skeletal muscle and tested whether enhancing NP signaling by preventing its clearance in skeletal muscle would improve exercise performance in mice. Under sedentary conditions, *Nprc* skeletal muscle knockout (MKO) mice showed comparable exercise performance to their floxed littermates in terms of maximal running velocity and total endurance running time. Eight weeks of voluntary running-wheel training in a young cohort increased exercise performance, but no significant differences were observed in MKO compared with control mice. Furthermore, 6-weeks of treadmill training in a relatively aged cohort also increased exercise performance compared with their baseline but did not result in an improvement in MKO mice compared with the controls. In summary, our study suggests that NP signaling is potentially important in myocytes but its function in skeletal muscle *in vivo* needs to be further studied in alternative physiological conditions or with new genetic mouse models.

## Introduction

Natriuretic peptides (NP) are hormones that are produced by the heart and endothelial cells, and they play essential roles in blood pressure control, cardiovascular homeostasis, and systemic metabolic regulation. There are three types of NP receptors (NPR), including the guanylyl cyclase-coupled receptors NPRA and NPRB, and the clearance receptor NPRC. Atrial natriuretic peptide (ANP) and B-type natriuretic peptide (BNP) bind to the NPRA receptor, while C-type natriuretic peptide (CNP) binds to the NPRB receptor. All three peptides bind with relatively equal affinity to NPRC, a transmembrane receptor that does not have a intracellular guanylyl cyclase domain (1). Binding of NP with NPRA and NPRB receptors activates their intracellular guanylyl cyclase activity to stimulate cyclic guanosine monophosphate (cGMP) production and increase cGMP-dependent protein kinase (PKG) phosphorylation of downstream targets. In contrast, binding with the NPRC receptor internalizes the peptide for degradation and thus effectively clears the peptide from circulation (2).

Exercise, a well-known modulator of bodyweight and systemic metabolism, can reduce the risk of developing obesity-related conditions such as Type 2 diabetes and cardiovascular disease (3). Rising obesity rates and accompanying co-morbidities in the population necessitate the ongoing search for ways to improve the beneficial metabolic effects of exercise on the human body. In healthy human subjects, prolonged endurance exercise has been shown to increase plasma NP levels (4). Previous reports have shown that NPs are inversely related to visceral fat, body mass index, and circulating insulin, and widely thought to influence obesity via their lipolytic effects in adipose tissue, fatty acid oxidation in skeletal muscle, and adipose tissue (5). This raised the question as to whether the metabolic benefit of exercise for obesity intervention could be mediated by the increases in plasma NP levels. In adipose tissues, NPs can increase adipose tissue ‘browning’ and mitochondrial biogenesis through PKG-mediated activation of peroxisome proliferator activated receptor gamma coactivator 1-α (PGC1α) and uncoupling protein 1 (UCP1) (6, 7). Skeletal muscle, given its relatively large percent of body mass, plays a crucial role in the regulation of exercise strength as well as stamina, and metabolic homeostasis. NP treatment increases mitochondrial metabolism, fat oxidation, and maximal respiration in human myotubes (8). In addition, *NPRA* (*NPR1*) gene expression is positively correlated to mRNA levels of *PGC1α* and several oxidative phosphorylation (OXPHOS) genes in human skeletal muscle, and the expression of *NPRA, PGC1α*, and OXPHOS genes is coordinately upregulated in response to aerobic exercise training in human skeletal muscle (8). These studies suggest a role for NP signaling in improving mitochondrial function and oxidative metabolism in muscle (8). Further studies of the metabolic function of NPs in skeletal muscle tissue could shed more light on their ability to combat obesity.

The NP clearance receptor NPRC plays essential roles in dictating NP signaling and its metabolic effect on target tissues. Whole-body and adipose tissue-specific deletions of the *Nprc* (*Npr3*) gene in mice result in reduced body fat, improved insulin sensitivity, elevated energy expenditure, and resistance to diet-induced obesity (6, 7). In those studies, we showed that mice with *Nprc* deletion in skeletal muscle are not protected against diet-induced obesity and insulin resistance (6) but the function of *NPRC* in skeletal muscle *per se* has not been well-studied. In the present work we sought to address the potential benefit of NPRC deficiency in skeletal muscle to determine whether it would improve the exercise performance and skeletal muscle metabolism.

## Results

### NP and NP receptor gene expression in C2C12 myocytes

To explore the role of natriuretic peptide signaling in regulating skeletal muscle function and metabolism, we first examined the expression of natriuretic peptide and receptor genes in C2C12 cells, a commonly used myocyte cell line. During the differentiation of C2C12 cells (**Fig. 1A**), the expression of NP receptor A (*Npra*, or *Npr1*) did not change. The expression of NP receptor B (*Nprb*, or *Npr2*) increased, while the expression of NP receptor C (*Nprc*, or *Npr3*) decreased (**Fig. 1B** and **Fig. S1**). Of note, the expression of CNP gene (*Nppc*), which acts on both NPRB and NPRC in an autocrine manner, is also induced during C2C12 cell differentiation (**Fig. S1**). The expression of the BNP gene (*Nppb*) increased in the first two-days of differentiation and then returned to the basal level, while the expression of ANP gene (*Nppa*) did not change during C2C12 expression (**Fig. S1**). In fully differentiated C2C12 myocytes, the expression of *Nprb* is the highest among the three NP receptors, and expression of *Nppc* is the highest among the three peptides (**Fig. 1B**). These data show that the natriuretic peptide and receptor genes are expressed and dynamically regulated during C2C12 myocyte differentiation.

**Figure 1.**
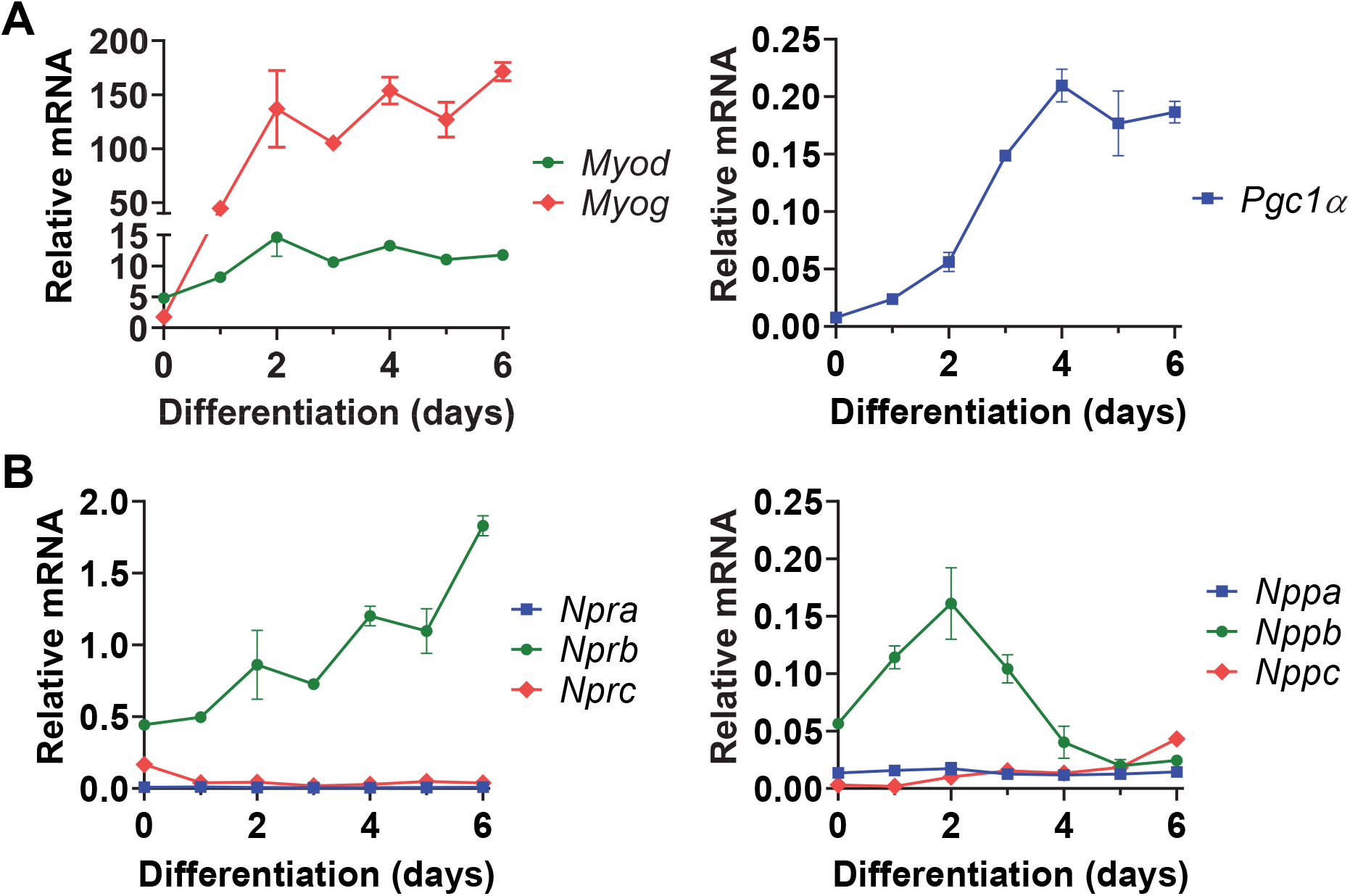
Expression of natriuretic peptide and receptor genes in C2C12 myocytes. mRNA levels of (A) myocyte marker genes *Myod, Myod* (left) and *Pgc1α* (right); (B) NP receptor genes *Npra, Nprb* and *Nprc* (left); and natriuretic peptide genes *Nppa, Nppb* and *Nppc* (right) in differentiating C2C12 myocytes. Data were normalized to internal control gene *36B4*.

### NPs stimulate PKG signaling in C2C12 myocytes

To further determine the effect of NPs on C2C12 myocytes, we next checked NP-stimulated cGMP signaling in C2C12 myocytes. Treatment with a membrane-permeable analog of cyclic guanosine monophosphate (cGMP), but not cyclic adenosine monophosphate (cAMP), increased the phosphorylation of Ser-239 (p-S^239^) in vasodilator-stimulated phosphoprotein (VASP), a known PKG substrate (**Fig. 2A**). Similarly, treatment with either ANP, BNP, or CNP increases p-S^239^ in VASP. The effect of CNP on p-S^239^ in VASP is the most robust among the three NPs (**Fig. 2B**), which is consistent with the observation that NPRB expression is the highest among the three NPRs in fully differentiated C2C12 myocytes (**Fig. 1B**). These data demonstrate that NPs can exert functional effects on C2C12 myocytes through the NPR-PKG signaling cascade.

**Figure 2.**
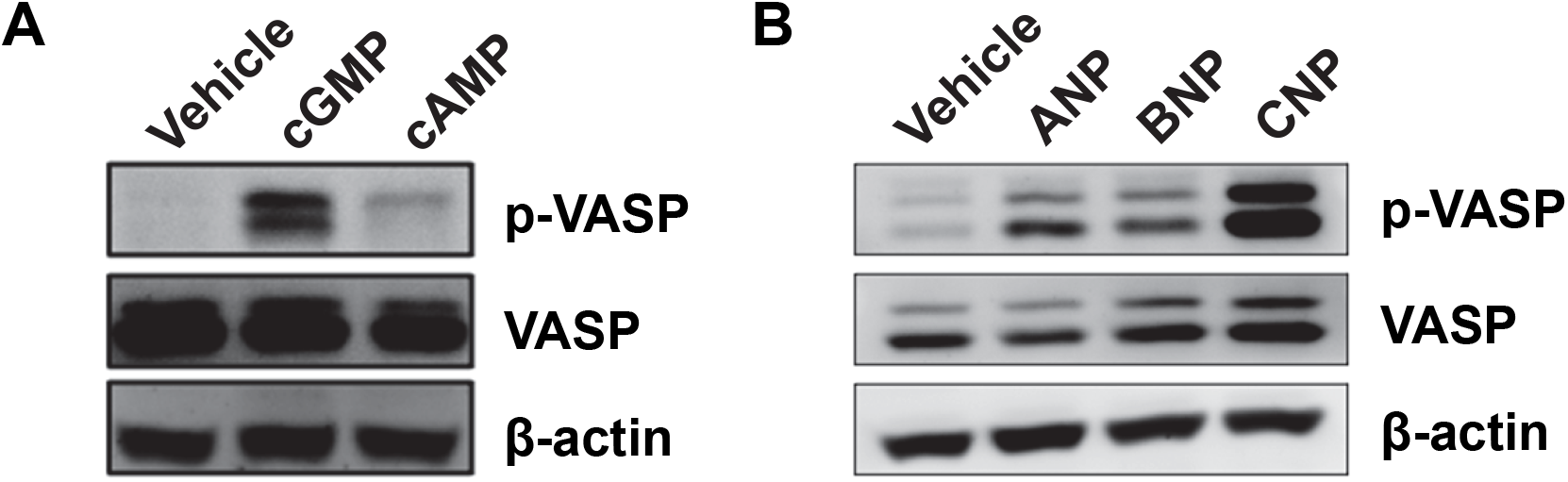
NPs stimulate PKG signaling in C2C12 myocytes. After six days of differentiation, C2C12 myocytes were serum-starved overnight and treated (A) with membrane-permeable 8-Br-cGMP (200nM) or 8-Br-cAMP (200nM); (B) with ANP (200 nM), BNP (200 nM) or CNP (200 nM) for 30 minutes. Levels of phosphorylated VASP (p-VASP), total VASP (VASP), and β-actin were then measured by Western blot analysis.

### Generation of *Nprc* skeletal muscle specific knockout (MKO) mice

Our data suggest that both NPRA and NPRB might be involved in the NP signaling in C2C12 myocytes. Since NPRC functions as a clearance receptor and all three NPs bind to NPRC, we next sought to generate a skeletal muscle specific *Nprc* knockout mice to determine the physiological effect of NP signaling in the skeletal muscle. We bred the *Nprc* floxed mice with *Myogenin*-Cre expressing mice to generate the skeletal muscle knockout (MKO) mice. As shown in our previous study (6), the expression of *Nprc* is relatively low in quadriceps femoris (QU), gastrocnemius (GA) and extensor digitorum longus (EDL) muscles, moderate in tibialis anterior (TA) muscle, and the highest in soleus muscle, while the expression of *Npra* is more consistent between these different types of muscles (**Fig. S2**) (6). In this regard, our gene expression analysis in the current manuscript will be primarily focused on the TA and soleus muscles. The *Nprc* MKO mice grow normally and do not display any gross morphological changes. As expected, the expression of *Nprc* was essentially absent in the TA and soleus muscles (**Fig. 3A**). The expression of *Npra, Nppa, and Nppc* did not change, but the expression of *Nprb* tended to increase in the skeletal muscle of *Nprc* MKO mice (**Fig. 3A, B**). The expression of *Pgc1α*, a transcriptional co-regulator involved in mitochondrial biogenesis, was comparable between genotypes in both TA and soleus muscles (**Fig. 3C**). Interestingly, the expression of *Uncoupling protein 3* (*Ucp3*), a gene linked to skeletal muscle oxidative metabolism and mitochondrial respiration, was significantly increased in both TA and soleus muscles (**Fig. 3C**).

**Figure 3.**
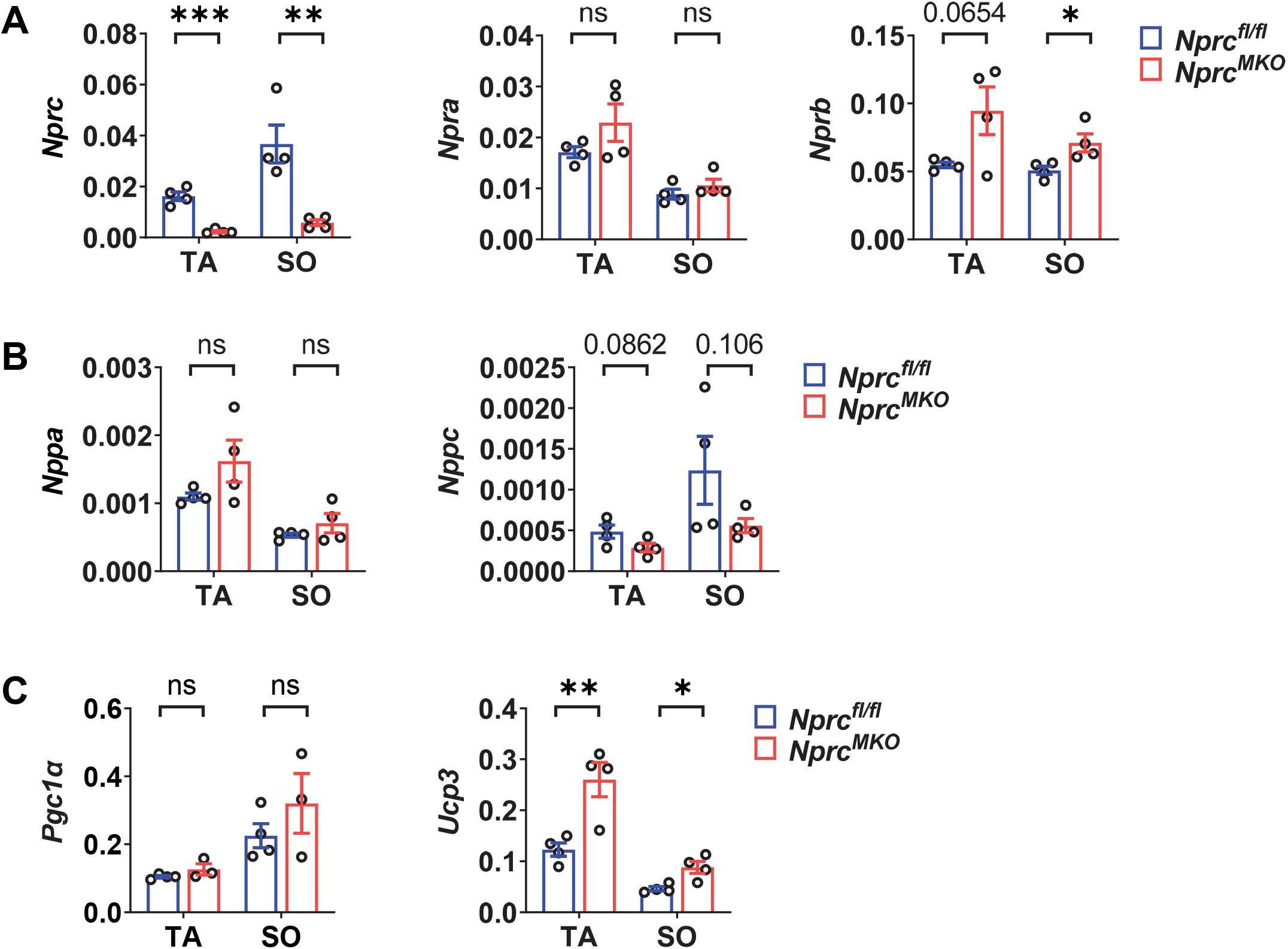
Gene expression profile in skeletal muscle of sedentary mice. (A) mRNA levels of (A) NP receptor genes *Nprc, Npra* and *Nprb*; (B) natriuretic peptide genes *Nppa* and *Nppc*; (*Nppb* was undetectable) and (C) *Pgc1α* and *Ucp3* in the tibialis anterior (TA) and solus (SO) muscles of control and *Nprc* MKO mice. Unpaired two-tailed Student’s t-tests, * p < 0.05; ** p < 0.01; *** p < 0.001; and ns: no statistical significance.

### Exercise performance of *Nprc* MKO after running-wheel training

Levels of cardiac natriuretic peptides have been reported to increase in healthy individuals after exercise training (4, 9-11). With a combination of increased systemic NP production and disrupted NP clearance in the skeletal muscle, we next sought to examine whether the metabolism and performance of *Nprc* MKO mice would be improved in the context of exercise training. We first checked the exercise performance of a sedentary cohort of 3- to 4-month-old male mice but did not observe any differences between genotypes regarding the maximal running speed and total endurance running time (**Fig. 4A, B**). We next set up a second cohort of mice for exercise training on voluntary running-wheels for eight weeks (**Fig. 5A**). Both *Nprc* MKO and control mice underwent similar amount of exercise training, as suggested by comparable running distance between two genotype group (**Fig. 5B**). Their exercise capacity after training was evaluated as shown in the sprint and endurance exercise tests. After the 8-week running-wheel training, the exercise performance of both genotypes was significantly improved compared to the sedentary mice (**Fig. 4**). However, both the maximal running speed and endurance running distance of *Nprc* MKO was not improved compared to that of the control mice (**Fig. 5C**).

**Figure 4.**
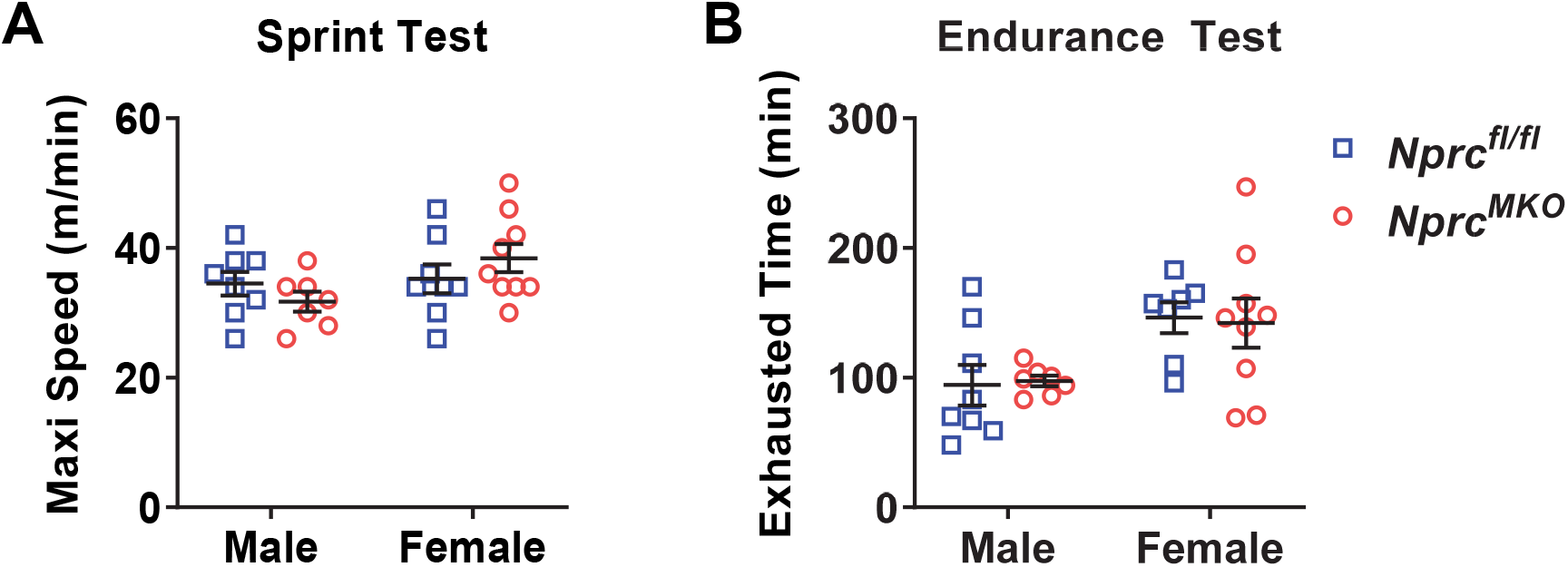
Exercise performance at sedentary status. A cohort of 3.5- to 5-month-old *Nprc* MKO and control mice were subjected to (A) sprint exercise test and (B) endurance exercise test to determine their maximal running speed and exhausted running time, respectively.

**Figure 5.**
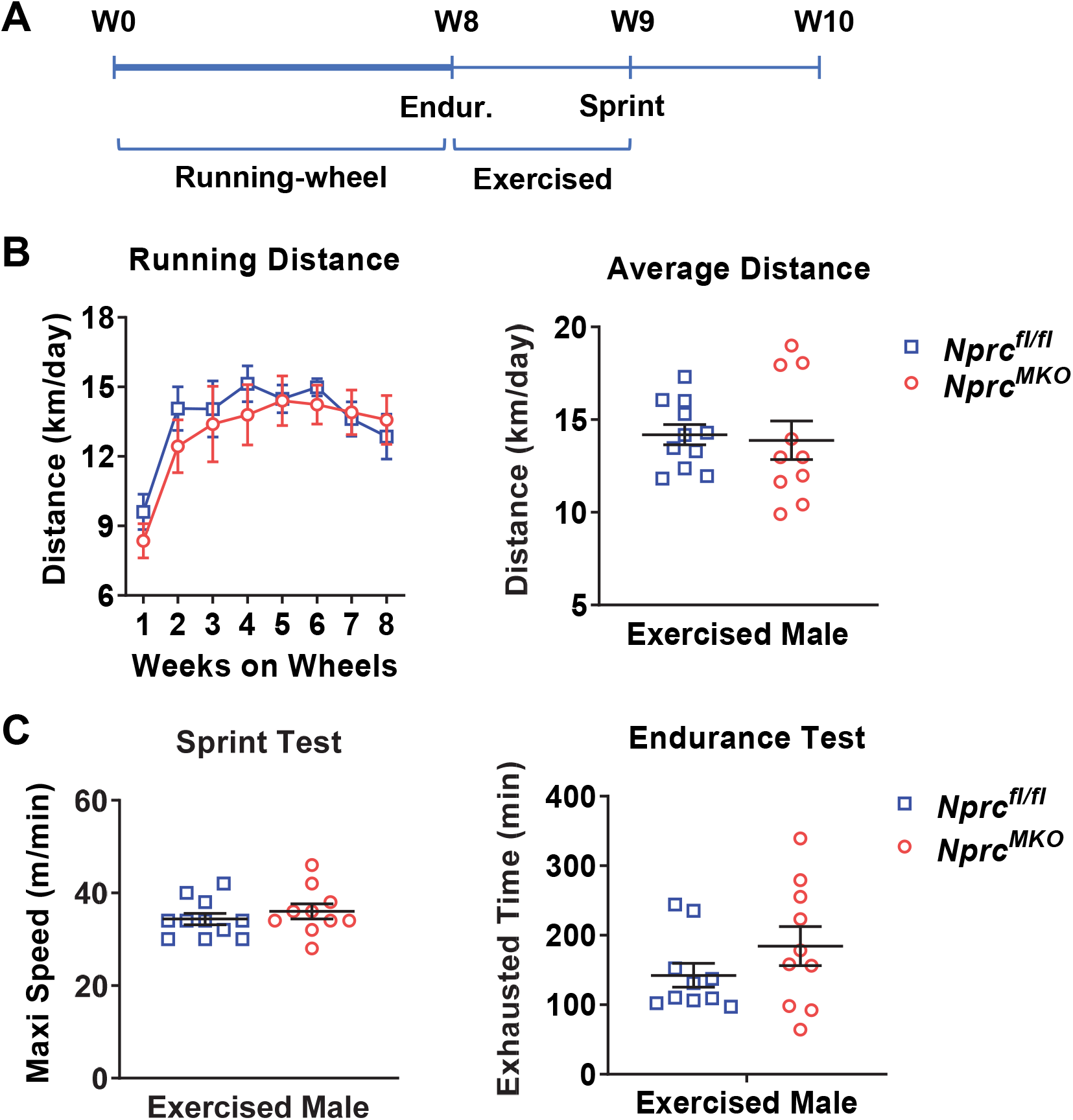
Exercise performance after 8 weeks of running-wheel training. (A) Timelines of running wheel training, endurance (Endur.) and sprint exercise tests. Six-week-old *Nprc* MKO and control male mice were trained in cages with a running-wheel outfitted with an odometer for eight weeks. (B) The daily running distances during running-wheel training. (C) After 8 weeks of training, mice were subjected to sprint exercise test (left) and endurance exercise test (right) to determine their maximal running speed and exhausted running time, respectively.

### Exercise performance of *Nprc* MKO after treadmill training

Previous studies have shown that exercise performance declines during the human life span and in aged mice (12, 13). Since we did not observe a robust improvement in the exercise capacity between *Nprc* MKO and control mice in relatively young cohorts, we next asked whether *Nprc* MKO mice would have better exercise capacity when older. A cohort of 1-year-old male and female mice were trained on treadmills for six weeks (**Fig. 6A**). In the males, six-week treadmill training greatly reduced their bodyweight and improved the exercise performance of both *Nprc* MKO and control mice, with increased maximal running speed in the stress test and longer running distance in the endurance test, compared to the baseline (**Fig. 6B-D**). In the females, six-week treadmill training also results in body weight loss and increased running distance in the endurance test, but the improvement in maximal running speed in the stress test was not as robust as that of the male mice (**Fig. 6B-D**). However, compared to the control mice, we did not observe any greater improvement in exercise performance in *Nprc* MKO mice (**Fig. 6B-D**). In addition, the gene expression profile was also comparable between *Nprc* MKO and control mice as well, including NP and NP receptors, *Pgc1α, Pgc1β, Ucp3*, Neprilysin (*Nep*) and Osteocrin (*Ostn*) (**Fig. 7**).

**Figure 6.**
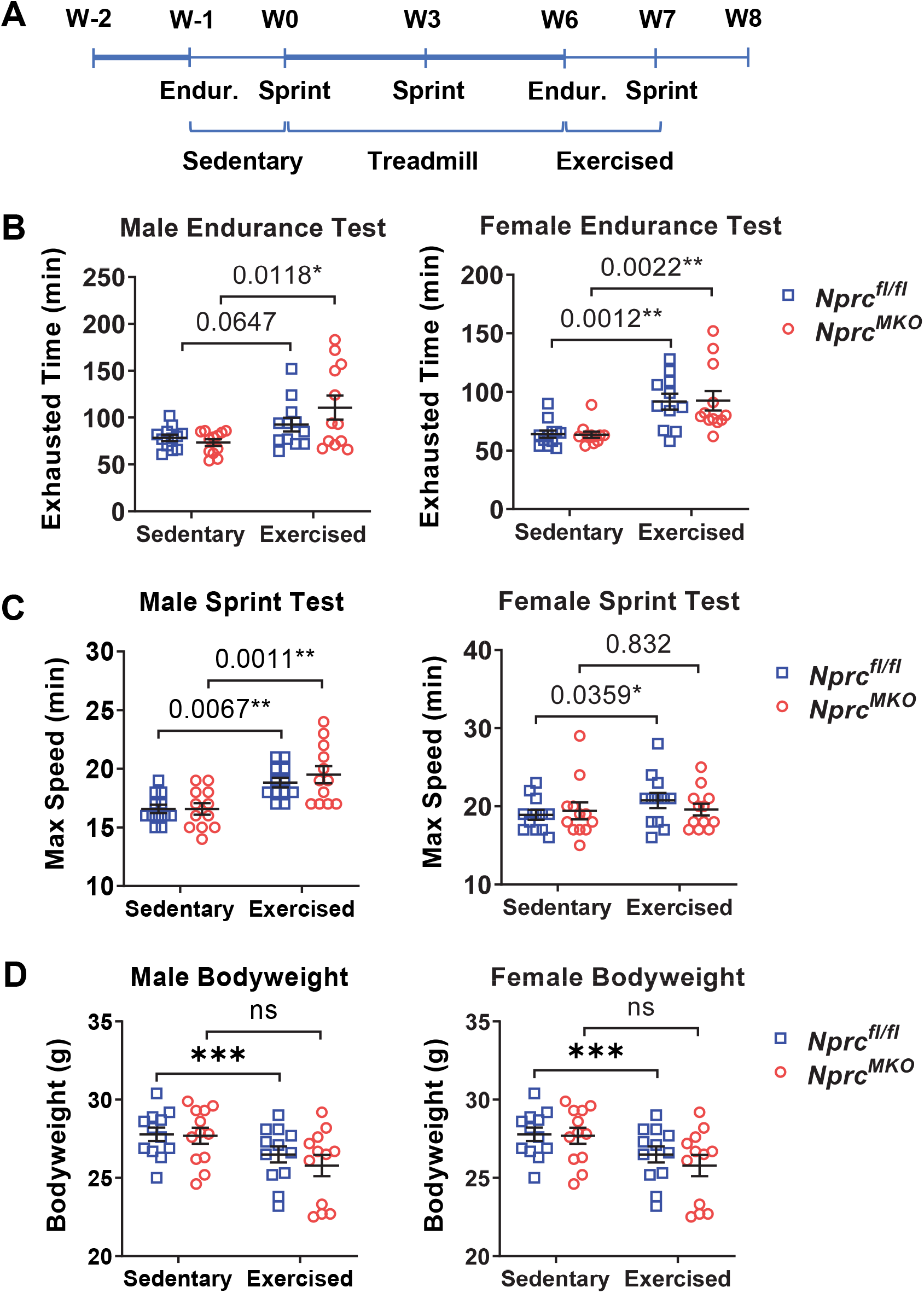
Exercise performance after 6 weeks of treadmill training. (A) Timelines of treadmill training, endurance (Endur.) and sprint exercise tests. Ten-month-old *Nprc* MKO and control male mice were trained on treadmill at a speed of 60% of their maximal running speed for six weeks. (B-C) Mice were subjected to (B) sprint exercise test and (C) endurance exercise test at baseline (Sedentary) and after treadmill training (Exercised) to determine their maximal running speed and exhausted running time, respectively. (D) The body weight of each group of mice at baseline and after treadmill training. Unpaired two-tailed Student’s t-tests, * p < 0.05; ** p < 0.01; *** p < 0.001; and ns: no statistical significance.

**Figure 7.**
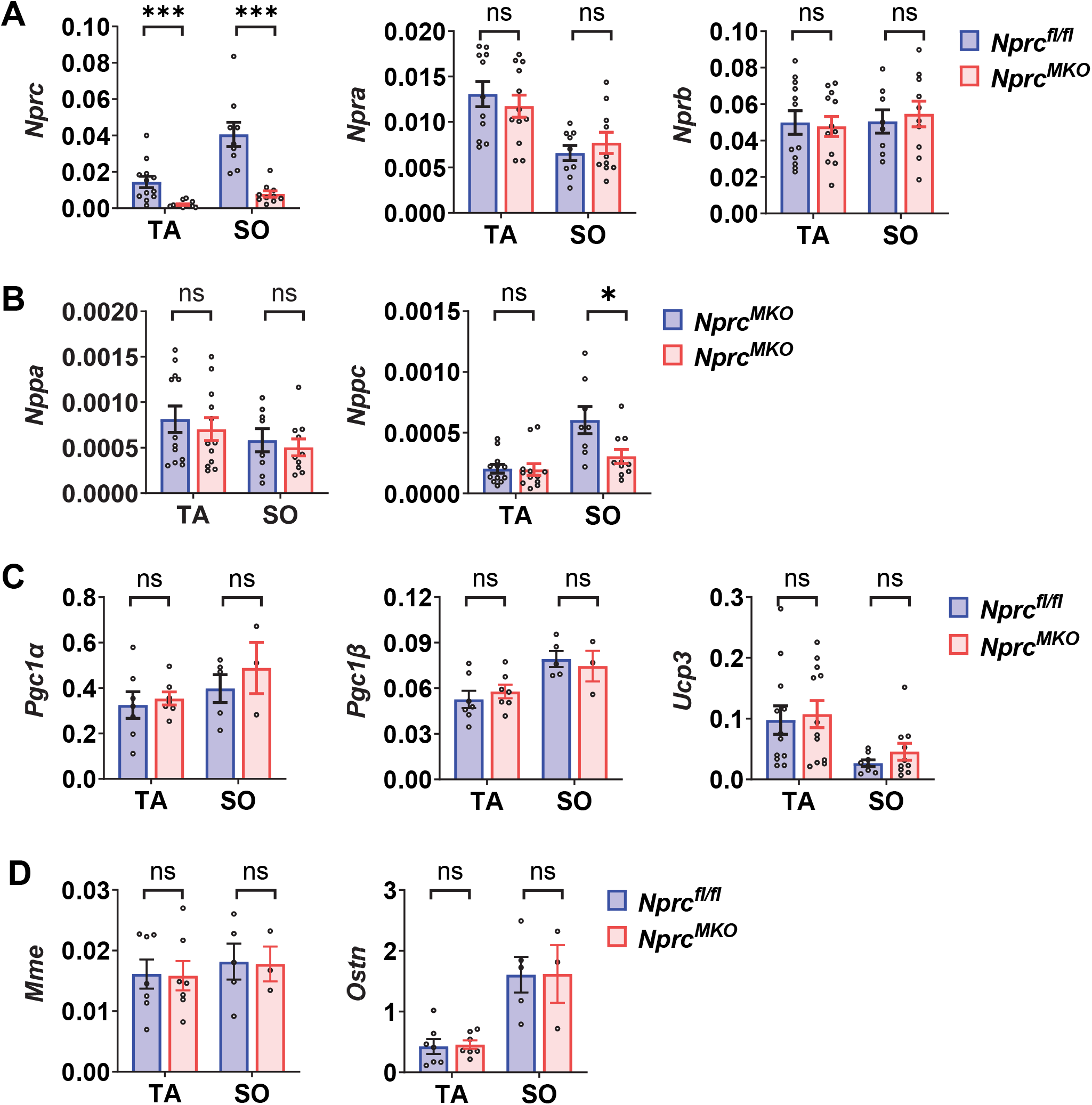
Gene expression profile in skeletal muscle of mice after treadmill training. (A) mRNA levels of (A) NP receptor genes *Nprc, Npra* and *Nprb*, (B) natriuretic peptide genes *Nppa* and *Nppc* (*Nppb* was undetectable), and (C) *Pgc1α, Pgc1β*, and *Ucp3*; and (D) neprilysin (*Mme*) and osteocrin (*Ostn*) in the tibialis anterior (TA) and solus (SO) of control and *Nprc* MKO mice. Unpaired two-tailed Student’s t-tests, * p < 0.05; *** p < 0.001; and ns: no statistical significance.

## Discussion

Evidence from human and rodent studies suggests that natriuretic peptide signaling has an important role in skeletal muscle metabolism. For example, chronic BNP infusion has been shown to increase glucose control, insulin sensitivity, and upregulation of lipid oxidative capacity in the skeletal muscle tissue of obese and Type 2 diabetic mice (14), and NPs have been reported to play an important role in the promotion of mitochondrial biogenesis and oxidative respiration in human skeletal muscle (8). The NPRC receptor is responsible for the vast majority of clearance and degradation of circulating NPs (2), thus we hypothesized that *Nprc* deletion would enhance NP signaling in the skeletal muscle for metabolic benefits and perhaps also result in improved exercise performance in the MKO group. However, our data did not show a clear improvement in the exercise performance of *Nprc* MKO mice regarding their endurance or sprint running capacity either under sedentary conditions or after exercise training, in both young and aged mice. This suggests that NPRC deletion in skeletal muscle might not exert a major physiological consequence for exercise capacity. The comparable exercise performance between *Nprc* MKO and floxed control mice suggests that a compensatory mechanism may exist to counteract the loss of NPRC, serving to maintain the homeostasis of NP signaling in skeletal muscles. Endogenous *Nprb* mRNA expression levels increase substantially in differentiated C2C12 myotubes (**Fig. 1**), suggesting a role for the CNP-NPRB pathway in regulating myocyte metabolism and function. In other studies (15), localized measurements of the gradient between arterial and veinous CNP concentrations across the femoral vein in humans found that musculoskeletal tissue made a measurable contribution to plasma CNP secretion levels in the body. Unlike ANP and BNP, CNP does not behave as a cardiac hormone and is present at extremely low levels in circulation, portraying CNP-NPRB signaling as a relatively localized pathway for NP action in skeletal muscle (16). Further study with loss of function mouse models, such as *Npra* or *Nprb* skeletal muscle knockout mice, will be necessary to clarify the contribution of different NP signaling pathways to skeletal muscle metabolism and exercise physiology in mice.

Proteolysis of NPs by neprilysin (NEP) typically acts in tandem with NPRC as an extracellular NP degradation agent (2). Administration of a NPRC-specific ligand C-ANF_4-23_ in rats has been shown to results in compensatory NEP activity, perhaps to lower ANP levels and maintain natriuretic peptide homeostasis (17). Since NEP and angiotensin converting enzyme inhibition have together improved morbidity and mortality in patients with heart failure by facilitating increased NP levels and binding to NP receptors (18), this NPRC-independent NP degradation pathway could function to compensate for the loss of NPRC to maintain the NP signaling in skeletal muscle. However at least at the gene expression level NEP was not different in the exercised mice. Further studies implementing NEP inhibition during treadmill training alongside NPRC deletion in the skeletal muscles might result in a larger improvement in exercise performance.

## Methods

### Reagents and antibodies

The membrane-permeable 8-Br-cGMP (8-bromoguanosine 3′,5′-cyclic monophosphate, B1381) and 8-Br-cAMP (8-bromoadenosine 3′,5′-cyclic monophosphate, B5386) were from Sigma. ANP (AS-20648), BNP (AS-24016), and CNP (AS-24244) were from AnaSpec. Antibodies used in this study included anti-p-Ser239 VASP (3114, Cell Signaling Technology), anti-total VASP (3112, Cell Signaling Technology), anti-GAPDH (sc25788, Santa Cruz; 10494-I-AP, ProteinTech), anti-rabbit IgG conjugated with alkaline phosphatase (A3687, Sigma).

### Cell lines

C2C12 mouse myoblast cells (ATCC, CRL-1772) were cultured in DMEM (4.5g/L of L-glucose, no sodium pyruvate) with 10% FBS, 1% penicillin-streptomycin. For differentiation, C2C12 cells were allowed to grow to confluency and differentiated in DMEM with 2% donor equine serum, 1µM insulin, 1% penicillin-streptomycin for 6 days.

### Mice

Mice with a floxed *Nprc* allele were generated as previously described (6). *Myogenin*-Cre mice were a gift from Eric Olson of University of Texas Southwestern Medical Center and bred with *Nprc* floxed mice to generate skeletal muscle specific knockout (MKO) mice. Mice were kept under a 12-hour light/12-hour dark cycle at constant temperature (23°C) with ad libitum access to water and mouse chow diet. All animal studies were approved by the Institutional Animal Care and Use Committee of Sanford Burnham Prebys Medical Discovery Institute and Vanderbilt University Medical Center in accordance with the National Institutes of Health (NIH) Guide for the Care and Use of Laboratory Animals.

### Exercise performance and training

#### Cohort 1: Sprint and endurance tests

Three- to 5-month-old *Nprc* MKO and control male mice were tested on an ECO-6M Treadmill (Columbus Instruments, Columbus, OH) with a 10 min warm-up at a speed of 10 m/min. For endurance test, the speed was then increased by 2 m/min every 5 min until reaching a final constant speed of 20 m/min. For sprint test, the speed was increased by 2 m/min every 2 min until the mice were exhausted. The point of exhaustion for both tests was reached when mice made continuous contact with the shock grid for 5 seconds.

#### Cohort 2: Running-wheel training

Six-weeks-old *Nprc* MKO and control male mice were individually housed in a cage with a running-wheel outfitted with an odometer for eight weeks. The daily running distance were recorded with the odometer. After 8-weeks training, mice were subjected to endurance and sprint tests as described in Cohort 1.

#### Cohort 3: Treadmill training

Ten-month-old *Nprc* MKO and control mice underwent a seven-day adaptation training procedure. For adaptation training, mice were placed on the treadmills for a total of 45 minutes, with an initial speed at 8m/min for the first 10 minutes and then with a constant speed at 10m/min afterwards. Mice that failed to complete at least five days of the adaptation training were removed from the study. After adaptation training, endurance and sprint tests were performed to determine the baseline performance. For endurance test, treadmill speed was initially set at 8 m/min and then increased by 1 m/min every 10 minute until exhaustion. For sprint test, treadmill speed was set at 8 m/min for 10 minutes and then increased by 1 m/min every 4 minutes until exhaustion. After baseline tests, mice were trained at 60% of their average maximum speeds for 1 h/day and 5 d/week for 3 weeks. At 3 weeks of training, mice underwent a second sprint test and trained at 60% of adjusted average maximum speeds for 1 h/day and 5 d/week for another 3 weeks. After 6 weeks of training in total, mice underwent final endurance and sprint tests to determine the final performance.

### RT-qPCR

Total RNA was extracted from C2C12 cells or tissues using TRIzol reagent (Invitrogen). C2C12 samples were then purified by isopropanol precipitation, and tissue samples by RNeasy Mini (Qiagen) or Quick-RNA Miniprep (Zymo Research) kits. cDNA was prepared with the High-Capacity cDNA Reverse Transcription (RT) kit (Applied Biology). Quantitative PCR (qPCR) were conducted with PowerUp SYBR Green Master Mix (Life Technologies) on the QuantStudio 6 Flex System (Applied Biosystems) according to the manufacturer’s protocols. The sequences of primers are listed in **Table S1**. RT-qPCR results were analyzed by ΔΔCt method, normalized to the reference gene, and expressed as mean ± SEM. *Ap3d1* and *36B4* were used as the reference genes for C2C12 cells and tissue samples, respectively,

### Western blot

Differentiated C2C12 cells were serum-starved for 48h in low-glucose Gibco DMEM and then stimulated with ANP, BNP, or CNP at 200nM for 30min, or subjected to a 30min treatment with 200nM 8-Br-cGMP as a positive control to induce downstream PKG activity and 200nM 8-Br-cAMP to act as a negative control. Cells were lysed and sonicated in lysis buffer (25 mM HEPES at pH 7.4, 150 mM NaCl, 5 mM EDTA, 5 mM EGTA, 5 mM glycerophosphate, 0.9% Triton X-100, 0.1% IGEPAL, 5 mM sodium pyrophosphate, 10% glycerol, 1x Complete protease inhibitor and 1x PhoSTOP phosphatase inhibitor cocktail). For Western blotting, 40 μg of protein was resolved by 10% SDS-polyacrylamide gel electrophoresis, transferred to nitrocellulose membranes (Bio-Rad), incubated overnight at 4 °C with primary antibodies, and followed by inoculation with alkaline phosphatase–conjugated secondary antibody for 1 hour at room temperature. Blots were incubated with Amersham ECF substrate (Cytiva, RPN5785) and images were acquired by Bio-Rad digital ChemiDoc MP with IR (VUMC Molecular Cell Biology Resource).

### Statistical analysis

GraphPad Prism 7 was used for statistical analysis. All data were presented as means ± SEM. Unpaired two-tailed Student’s t-tests were used to determine the differences between groups. Statistical significance was defined as p-value < 0.05.

## Supporting information

Nprc muscle supplement

## Acknowledgment

We thank Wei Zhang and Huafeng Fang for excellent technical assistance and animal husbandry. We thank Ling Lai, Meghan Gabriel, and Felipe Castellani Gomes Dos Reis for advice on mouse exercise training protocols. This work was supported by NIH R01 DK116625 (SC).

## References

1. Anand-Srivastava MB, and Trachte GJ. Atrial natriuretic factor receptors and signal transduction mechanisms. Pharmacological reviews 45: 455–497, 1993.

2. Potter LR. Natriuretic peptide metabolism, clearance and degradation. The FEBS journal 278: 1808–1817, 2011.

3. Melmer A, Kempf P, and Laimer M. The role of physical exercise in obesity and diabetes. Praxis (Bern 1994) 107: 971–976, 2018.

4. Hamasaki H. The effects of exercise on natriuretic peptides in individuals without heart failure. Sports (Basel) 4: 2016.

5. Gruden G, Landi A, and Bruno G. Natriuretic peptides, heart, and adipose tissue: new findings and future developments for diabetes research. Diabetes care 37: 2899–2908, 2014.

6. Wu W, Shi F, Liu D, Ceddia RP, Gaffin R, Wei W, Fang H, Lewandowski ED, and Collins S. Enhancing natriuretic peptide signaling in adipose tissue, but not in muscle, protects against diet-induced obesity and insulin resistance. Science signaling 10: 2017.

7. Bordicchia M, Liu D, Amri EZ, Ailhaud G, Dessi-Fulgheri P, Zhang C, Takahashi N, Sarzani R, and Collins S. Cardiac natriuretic peptides act via p38 MAPK to induce the brown fat thermogenic program in mouse and human adipocytes. The Journal of clinical investigation 122: 1022–1036, 2012.

8. Engeli S, Birkenfeld AL, Badin PM, Bourlier V, Louche K, Viguerie N, Thalamas C, Montastier E, Larrouy D, Harant I, de Glisezinski I, Lieske S, Reinke J, Beckmann B, Langin D, Jordan J, and Moro C. Natriuretic peptides enhance the oxidative capacity of human skeletal muscle. The Journal of clinical investigation 122: 4675–4679, 2012.

9. Ohba H, Takada H, Musha H, Nagashima J, Mori N, Awaya T, Omiya K, and Murayama M. Effects of prolonged strenuous exercise on plasma levels of atrial natriuretic peptide and brain natriuretic peptide in healthy men. American heart journal 141: 751–758, 2001.

10. Clarkson PB, Wheeldon NM, MacFadyen RJ, Pringle SD, and MacDonald TM. Effects of brain natriuretic peptide on exercise hemodynamics and neurohormones in isolated diastolic heart failure. Circulation 93: 2037–2042, 1996.

11. Krupicka J, Janota T, Kasalová Z, and Hradec J. Effect of short-term maximal exercise on BNP plasma levels in healthy individuals. Physiological research / Academia Scientiarum Bohemoslovaca 59: 625–628, 2010.

12. Ganse B, Ganse U, Dahl J, and Degens H. Linear decrease in athletic performance during the human life span. Frontiers in physiology 9: 1100, 2018.

13. Haramizu S, Ota N, Hase T, and Murase T. Aging-associated changes in physical performance and energy metabolism in the senescence-accelerated mouse. The journals of gerontology Series A, Biological sciences and medical sciences 66: 646–655, 2011.

14. Coue M, Badin PM, Vila IK, Laurens C, Louche K, Marques MA, Bourlier V, Mouisel E, Tavernier G, Rustan AC, Galgani JE, Joanisse DR, Smith SR, Langin D, and Moro C. Defective natriuretic peptide receptor signaling in skeletal muscle links obesity to type 2 diabetes. Diabetes 64: 4033–4045, 2015.

15. Palmer SC, Prickett TC, Espiner EA, Yandle TG, and Richards AM. Regional release and clearance of C-type natriuretic peptides in the human circulation and relation to cardiac function. Hypertension 54: 612–618, 2009.

16. Pandey KN. Emerging roles of natriuretic peptides and their receptors in pathophysiology of hypertension and cardiovascular regulation. J Am Soc Hypertens 2: 210–226, 2008.

17. Hashimoto Y, Nakao K, Hama N, Imura H, Mori S, Yamaguchi M, Yasuhara M, and Hori R. Clearance mechanisms of atrial and brain natriuretic peptides in rats. Pharmaceutical research 11: 60–64, 1994.

18. Jhund PS, and McMurray JJ. The neprilysin pathway in heart failure: a review and guide on the use of sacubitril/valsartan. Heart 102: 1342–1347, 2016.

