## Supplementary material for "Exercise performance is not improved in mice with skeletal muscle deletion of natriuretic peptide clearance receptor": Nprc muscle supplement

**Supplementary Data**

**Figure S1. Expression of natriuretic peptide and receptor genes in C2C12 myocytes.**

Fold change of the mRNA levels of (A) NP receptor genes *Npra*, *Nprb* and *Nprc*; and (B) natriuretic peptide genes *Nppa*, *Nppb* and *Nppc* in differentiating C2C12 myocytes. Data were normalized to baseline at day 0 as fold change.

**Figure S2. Expression of *Nprc* and *Npra* in skeletal muscles.**

mRNA levels of (A) *Nprc* and (B) *Npra* in quadriceps (QU), gastrocnemius (GA), tibialis anterior (TA), soleus (SO), extensor digitorum longus (EDL) muscles of *Nprc* MKO and control mice (n=4-5). Numbers are the average relative Ct values of the control mice, and the average Ct value of internal control gene *36B4* is set as 18. (Wu and Shi et al. *Science Signaling*, 2017)

**Table S1. Sequences of QPCR primers.**

| **Genes** | **Fwd (5'-3')** | **Rev (5'-3')** |
|---|---|---|
| *Npra* | TGGAGACACAGTCAACACAGC | CGAAGACAAGTGGATCCTGAG |
| *Nprb* | GAAGGCCTGGACCTCAGTC | TCAGTTGTGTCCGGTCAATG |
| *Nprc* | AGCTGGCTACAGCAAGAAGG | CGGCGATACCTTCAAATGTC |
| *Nppa* | GCTTCCAGGCCATATTGGAG | GGGGGCATGACCTCATCTT |
| *Nppb* | CCAGCAGAGACCTCAAAATTCC | AACTTCAGTGCGTTACAGCC |
| *Nppc* | TGAGCGGTCTGGGATGTTAG | ATTGCGTTGGAGGTGTTTCC |
| *Myod* | AGCACTACAGTGGCGACTCA | GGCCGCTGTAATCCATCAT |
| *Myog* | CCTTGCTCAGCTCCCTCA | TGGGAGTTGCATTCACTGG |
| *Pgc1α* | CGGAAATCATATCCAACCAG | TGAGAACCGCTAGCAAGTTTG |
| *Pgc1β* | CTCCAGTTCCGGCTCCTC | CCCTCTGCTCTCACGTCTG |
| *Ucp1* | GGCCTCTACGACTCAGTCCA | TAAGCCGGCTGAGATCTTGT |
| *Neprilysin* | GCCAAAGCAAGCAGCTAAAG | CTGATTTCGGCCTGAGGAATAA |
| *Osteocrin* | CCATGGATCGGATTGGTAGA | TCTGTGCCATCTCACACAAGT |

**FigS1**

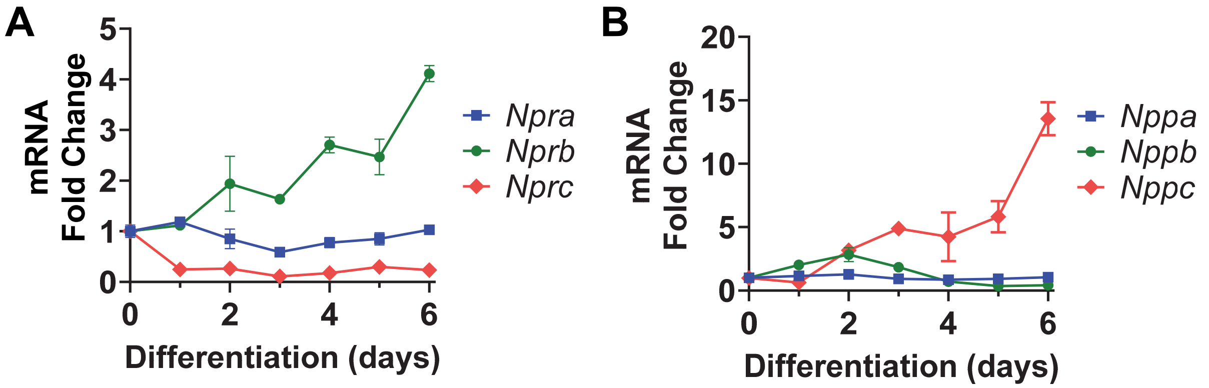


**Figure S1. Expression of natriuretic peptide and receptor genes in C2C12 myocytes.** Fold change of the mRNA levels of (A) NP receptor genes *Npra*, *Nprb* and *Nprc*; and (B) natriuretic peptide genes *Nppa*, *Nppb* and *Nppc* in differentiating C2C12 myocytes. Data were normalized to baseline at day 0 as fold change.

FigS2

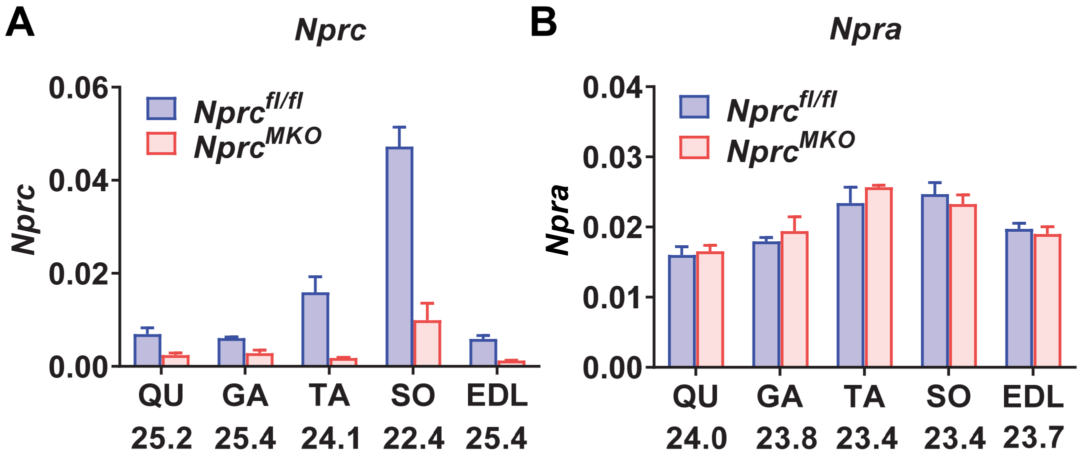


**Figure S2. Expression of *Nprc* and *Npra* in skeletal muscles.** mRNA levels of (A) *Nprc* and (B) *Npra* in quadriceps (QU), gastrocnemius (GA), tibialis anterior (TA), soleus (SO), extensor digitorum longus (EDL) muscles of *Nprc* MKO and control mice (n=4-5). Numbers are the average relative Ct values of the control mice, and the average Ct value of internal control gene *36B4* is set as 18. (Wu and Shi et al. *Science Signaling*, 2017)
